# Norm-like vocal behavior in a nonhuman primate

**DOI:** 10.64898/2026.08.03.742499

**Authors:** Nikhil Phaniraj, Monika Mircheva, Judith M. Burkart

**Affiliations:** Institute of Evolutionary Anthropology, University of Zurich; Winterthurerstrasse 190, 8057 Zürich, Switzerland; Neuroscience Center Zurich (ZNZ), University of Zurich and ETH Zurich; Winterthurerstrasse 190, 8057 Zürich, Switzerland; Center for the Interdisciplinary Study of Language Evolution (ISLE), University of Zurich; Affolternstrasse 56, 8050 Zürich, Switzerland

## Abstract

Human interlocutors have strong expectations that guide when, how, and what is communicated, but the evolutionary origins of such normative expectations remain poorly understood. Vocal accommodation in marmosets, where individuals converge their calls toward their partners’, offers a model for investigating their evolutionary roots. Using a fully automated closed-loop interactive playback paradigm, we presented marmosets with partner calls that converged toward or diverged from their own call structure. Marmosets responded affiliatively to converging calls, but with protest vocalizations to diverging calls. Importantly, the protests ceased when calls stopped diverging. Responses were partner-specific, indicating that marmosets expect bonded partners, but not strangers, to converge during a conversation. Moreover, individuals modified their calls to restore similarity to dissimilar partner, but not stranger playbacks. These findings identify marmoset vocal accommodation as socially regulated behavior with normative dimensions, suggesting that communicative expectations may have predated language and scaffolded the emergence of human linguistic conventions.

## Main

Human language is a socially regulated system in which communication is guided by shared expectations about how and when signals are exchanged^1–5^. These expectations constitute a complex framework of social norms that shape the timing, manner, and content of communication. Social norms are defined by four core features: (i) they are expressed *consistently* across individuals at the level in which the norm operates (e.g., individuals of a social group or a species); (ii) they elicit *conformity* with few violations; (iii) they serve a *functional role* in social life, and (iv) they are maintained through *social reinforcement* mechanisms such as approval or disapproval^6–9^.

The evolutionary origins of communicative social norms remain unclear. Although great apes show gestural communication where group members share conventions for how certain gestures are used and interpreted, evidence for a normative organization in their communication is absent^10–13^. In particular, evidence for approval or disapproval of appropriate use is critically lacking in apes and other nonhuman animals^14^. As a result, little is known about whether and how normativity may have emerged before the evolution of human language.

Vocal accommodation in the common marmoset (*Callithrix jacchus*) offers a promising window into this question. In this species, individuals modify the acoustic structure of their calls so that they become more similar to those of their social partners^15^ (i.e., convergent accommodation, henceforth ‘convergence’). Prior work shows that this phenomenon is expressed *consistently* across individuals^15,16^, that convergence is the dominant pattern with only rare cases of divergent accommodation (involving increased dissimilarity to a social partner’s calls, henceforth ‘divergence’)^16^, suggesting *conformity*, and that it has a *functional role* in the formation and maintenance of social bonds^17,18^. These features align with the first three criteria for social norms, yet one critical component remains untested: whether vocal accommodation is shaped by *social reinforcement*.

To test whether marmosets socially reinforce vocal accommodation, we employed synthetically generated calls in a fully automated closed-loop playback paradigm (Fig 1A) that leveraged the natural temporal dynamics of turn-taking to simulate vocal convergence toward or divergence away from a focal individual (Fig. 1B, C). We studied 24 adult marmosets constituting 12 male-female breeding pairs. Each pair was first habituated to a sound-proof experimental room in a separate study, where pair members were separated by a visual barrier and produced spontaneous affiliative^19^ phee calls in an antiphonal setting^20^ (Fig. 1A). This acclimation ensured that individuals expected to hear their partner during the experiments. We identified 1218 high-quality phee calls recorded from these sessions, which served as the acoustic corpus for playback and synthetic stimuli.

**Fig 1.**
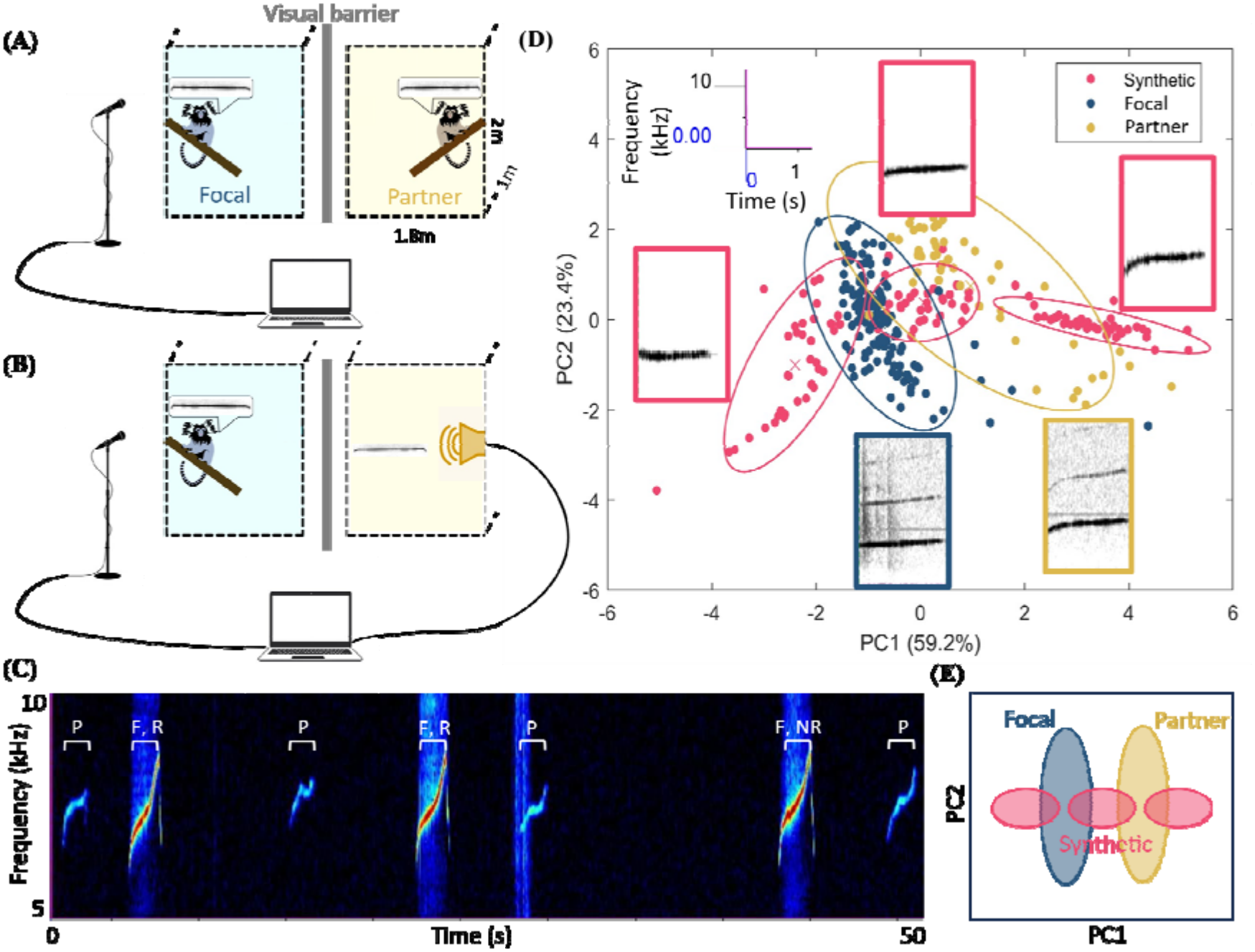
Interactive closed-loop acoustic playback experiments. **(A)** Habituation setup. Pair-bonded marmosets were placed in separate enclosures divided by an opaque visual barrier and communicated naturally through antiphonal phee calling. These sessions established an expectation that the partner was located behind the barrier and provided the natural call corpus used for stimulus generation. **(B)** Experimental setup. During playback experiments, the partner was replaced by a speaker controlled by a real-time laptop-based playback system. The system detected phee calls produced by the focal individual and generated interactive playback responses with naturalistic antiphonal timing. **(C)** Example segment of a closed-loop interaction between a focal marmoset and the playback system. Spectrograms show alternating phee call produced by the playback system and the focal individual. Calls labeled “R” were classified a valid antiphonal responses, whereas the call labeled “NR” was excluded because it occurred more than 12.4 s after the preceding playback. Abbreviations: P, playback call; F, focal call; R, response; NR, non-response. **(D)** Acoustic space of a subset of natural and synthetic phee calls from one representative pair. Blue points represent calls from the focal individual, yellow points represent calls from the partner, and pink points represent synthetic calls generated by morphing between and beyond the vocal structures of the pair. Calls are visualized in the space defined by the first two principal components of six acoustic features. Ellipses indicate 95% confidenc intervals for each call distribution and are used for visualization purposes only. Representative spectrograms from each call category are shown with a common time and frequency scale displayed on the top left. For each dyad, 60 calls per cluster were synthesized. **(E)** A highly simplified visualization of the clusters formed by focals’ (blue), partners’ (yellow) and synthetic (pink) calls in an arbitrary acoustic space.

To systematically manipulate vocal similarity, we generated synthetic phee calls spanning the acoustic space between and beyond each pair’s calls (Fig.1D, E). For every breeding pair, we synthesized 60 calls intermediate between the partners, 60 more dissimilar to the female than the male’s natural calls, and 60 more dissimilar to the male than the female’s, yielding 180 synthetic calls per pair and 2160 across all 12 pairs. Calls were synthesized by extracting amplitude and fundamental frequency trajectories from natural calls and representing them in a shared acoustic space. By assigning different weights to each partner’s call, we generated stimuli that varied systematically in similarity. For example, weighing a stimulus toward one partner produced a call resembling the call of that individual, whereas weighing away from a partner produced a call that was increasingly dissimilar. Together with the 1218 natural phee recordings, these stimuli formed an acoustic corpus of 3378 calls used for playback experiments. Across all conditions, we recorded and analyzed 1409 phee calls from focal individuals in response to the playbacks, allowing us to examine changes in interaction dynamics and acoustic structure of the responses.

In experiment 1, eight breeding pairs (n=16) participated in three 20-minute stimulus conditions presented on separate days in a randomized order (Fig. 2C). Animals heard either playbacks of partners’ natural phees (partner control: P-ctr), partner-like synthetic phees that progressively converged toward the focal individual’s own phees (partner-convergence-similar calls: P-con-sim) or partner-like synthetic phees that diverged from the focal’s phees (partner-divergence-dissimilar calls: P-div-dis). A session started with a 2-minute playback of natural partner phees that constituted the pre-treatment baseline, followed by 16 minutes of natural partner phees (P-ctr) or convergence (P-con-sim) or divergence (P-div-dis), and ended with another 2-minute post-treatment baseline of natural partner phees. All stimuli were delivered in an automated interactive closed-loop format, triggering playback in response to each phee produced by the focal animal, with a latency distribution modelled on natural antiphonal calling^21^ (Fig.1B, 1D, 2A, Movie S1). This design allowed us to examine how marmosets respond to changes in vocal similarity during interactional phee call exchange sequences, and whether such changes elicit affiliative or agonistic responses.

**Fig 2.**
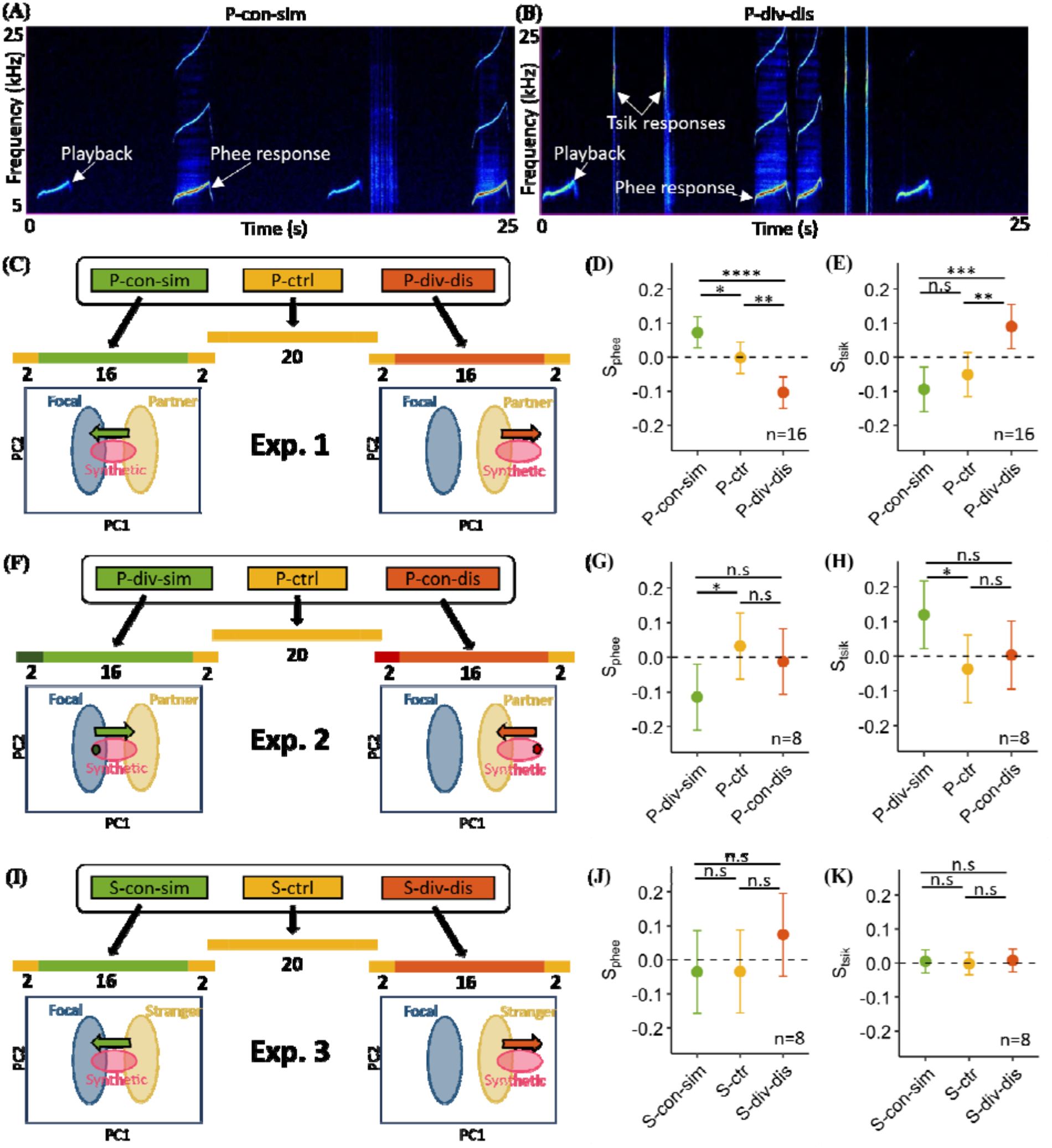
Marmosets respond affiliatively to vocal convergence and aversively to vocal divergence. **(A)** Representative spectrogram segment from a simulated convergence condition (P-con-sim) showing antiphonal phee responses by the focal individual to playback calls that progressively converged toward its own vocal structure. **(B)** Representative spectrogram segment from a simulated divergence condition (P-div-dis) showing both phee responses and aversive tsik vocalizations produced in response to playback calls that progressively diverged away from the focal individual’s calls. **(C, F, I)** Schematic representations of the playback conditions used in experiments 1, 2, and 3, respectively. Colored bars illustrate the temporal progression of playback stimuli across the 20-minute sessions, with numbers showing the duration of the subsections, and arrows within acoustic space diagrams indicate the direction of change in vocal similarity relative to the focal individual. **(C)** Experiment 1 tested responses to partner-derived playbacks. In the partner control condition (P-ctrl), subjects heard natural calls from their partner throughout the session. In the convergence condition (P-con-sim), partner-like synthetic calls gradually shifted toward the focal individual’s own vocal structure. In the divergence condition (P-div-dis), partner-like synthetic calls gradually shifted away from the focal individual’s calls. **(F)** Experiment 2 tested whether responses depended on the direction of acoustic change rather than absolute similarity. In the reversed divergence condition (P-div-sim), calls initially highly similar to the focal (dark green circle) gradually shifted toward the partner’s natural calls. In the reversed convergence condition (P-con-dis), calls initially highly dissimilar to the focal (red circle) gradually shifted toward the partner’s calls. **(I)** Experiment 3 tested partner specificity using unfamiliar individuals. Subjects experienced natural stranger calls (S-ctrl), stranger-like synthetic calls converging toward the focal individual’s calls (S-con-sim), and stranger-like synthetic calls diverging away from them (S-div-dis). **(D, E, G, H, J, K)** Model estimated phee response scores (S_phee_; **D, G, J**) and tsik response scores (S_tsik_; **E, H, K**) for experiments 1, 2, and 3, respectively. Points indicate estimated marginal means and error bars denote 95% confidence intervals. Positive scores indicate elevated responsiveness relative to the expected baseline trajectory, whereas negative values indicate reduced responsiveness. Sample sizes are shown within panels. Abbreviations: n.s., not significant; *p < 0.05; **p < 0.01; ***p < 0.001; ****p < 0.0001.

If vocal accommodation functions as a socially maintained norm, then marmosets should show affiliation towards partners that converge toward them and aversion towards those that diverge. We therefore predicted more affiliative phee responses during simulated convergence (P-con-sim) and fewer during simulated divergence (P-div-dis) relative to natural partner interactions. In addition to phees, we also monitored the production of tsik calls. Tsiks are typically emitted in negative or high-arousal situations, including alarm, mobbing, and social conflict^22,23^. They are also produced in the cases of frustation (for example, tsiks are directed at humans when they move a highly desired food item out of reach of the subjects). We therefore treated tsik production as an indicator of aversive reactions (“protests”) to the playback stimuli and a potential mechanism of negative social reinforcement. Under the social-norm hypothesis, we predicted elevated tsik responses to simulated divergence (P-div-dis) and no difference between tsik responses to simulated convergence (P-con-sim) and P-ctr.

Previous studies show that marmoset calling rates vary across sessions and typically decline exponentially within sessions^24–26^. We therefore quantified responses relative to session-specific expectations rather than call counts (Fig. S1). For each trial, mean response probabilities during the first and final two minutes served as pre- and post-baselines, and an expected exponential decay function was fit across the intervening period. A call was considered a response if it occurred within 12.4s of the playback^21^. We then calculated a response score (S_call_) for each call type by integrating deviations between observed calling probability and the expected trajectory using a moving two-minute window. A response score of S_call_ translates to 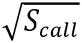 greater magnitude of probability to respond to the playbacks compared to the “expected” probability of response for 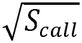 fraction of the session. Positive values indicate higher-than-expected responsiveness and negative values indicate suppression, with scores ranging from -1 to 1.

## Results

### Marmosets preferentially respond to convergence with affiliative calls

Linear mixed effects modeling identified stimulus condition as a strong predictor of phee responses (type III ANOVA, F_2,32_ = 21.388, p = 1.27e-06). Adding age, sex, presentation order, or their interactions with condition did not significantly improve the model fit (all likelihood ratio tests p > 0.1, Table S1). Response scores in the partner control condition did not differ from zero, indicating that responses to natural partner calls indeed closely followed the expected decay function, thus validating the baseline approach (Fig. 2D).

Marmosets showed significantly elevated phee responses (S_phee_) when exposed to partner-like synthetic calls that gradually converged toward their own vocal structure, compared to P-ctr, suggesting a highly affiliative reaction^19^, which could act as positive social reinforcement (Fig. 2D, pairwise comparisons in LMM with Tukey adjustment, n=16, df = 34.1, t ratio = 2.665, p = 0.0306). In contrast, S_phee_ were significantly reduced compared to P-ctr when calls gradually diverged away from the focal individual’s calls, which could act as negative social reinforcement (Fig. 2E, pairwise comparisons in LMM with Tukey adjustment, n= 16, df = 34.1, t ratio = 3.642, p = 0.0025). Marmosets were thus more likely to reply with phee calls relative to control to converging playbacks and less likely to diverging playbacks.

### Marmosets respond to divergence with aversive calls

In addition to antiphonal phee calls, focal individuals also responded to playback stimuli with tsik vocalizations (Fig. 2B). We quantified tsik responses (S_tsik_) using the same session-normalized procedure as applied to phee calls. Linear mixed effects modeling identified playback condition as a significant predictor of response score (type III ANOVA, F_2,32_ = 12.352, p = 1.06e-04) and adding age, sex, or presentation order did not improve model fit (all likelihood ratio tests p > 0.5, Table S2). Marmosets showed significantly elevated S_tsik_ during playback sequences in which calls gradually diverged away from the focal individual’s own vocal structure, compared to P-ctr, suggesting negative social reinforcement (Fig. 2E, pairwise comparisons in LMM with Tukey adjustment, n = 16, df = 34.1, t ratio = -3.519, p = 0.0035, Movie S2). The convergence condition, however, did not significantly differ from P-ctr. Moreover, marmosets stopped producing elevated tsik responses when playbacks reverted to the partner’s natural calls during the post-treatment baseline (Fig. S2). Marmosets thus showed elevated tsik responses to a diverging sequence of playbacks which stopped as soon as the calls returned to baseline, but not to a converging playback sequence. This is consistent with negative social reinforcement in response to a diverging partner, which stops when a putative effect on the partner (i.e., reverting to normal phees) is achieved.

### Responses depend on the direction of vocal change, not absolute similarity

The increased tsik responses during divergence playbacks could reflect three processes. First, marmosets may have reacted to the synthetic calls themselves, eliciting tsik responses to artificially sounding calls. Second, marmosets may simply have responded negatively to calls that were highly dissimilar to their own calls compared to their partner’s natural calls, regardless of whether they were synthetic. Third, marmosets may have reacted to calls becoming progressively less similar to their own over time, indicating sensitivity to the direction of change rather than absolute dissimilarity. To distinguish between these possibilities, we conducted experiment 2 in which the same synthetic stimuli were presented but in reverse order.

We tested four breeding pairs (n=8, based on the power analysis of the results from experiment 1) across three 20-minute playback conditions presented on separate days in randomized order (Fig. 2F). Two pairs had previously undergone experiment 1 and two were naïve. As before, animals heard either natural partner calls (P-ctr) or partner-like synthetic calls in a closed-loop antiphonal playback paradigm. In the test conditions of experiment 2, we used the same calls as in the test conditions of experiment 1, but reversed the stimulus order (Fig. 2F).

The divergence condition began with calls highly similar to the focal individual (2-minute pre-baseline), which then shifted gradually toward the partner’s natural phees (16-minute reversed convergence: P-div-sim), followed by a 2-minute post-baseline of natural partner phees. If the tsiks were simply a response to the calls being highly dissimilar to their own calls compared to the normal calls of the partner, then marmosets should not respond with tsiks in this condition. Only if the marmosets were sensitive to the direction of change (divergence) instead of absolute dissimilarity, would we expect greater tsik responses. The convergence condition began with calls highly dissimilar to the focal individual (2-minute pre-baseline) and then gradually shifted toward the partner’s natural phees (reversed divergence: P-con-dis). If marmosets reacted specifically to divergence, we would not expect elevated tsik responses. If marmosets instead responded to highly dissimilar or synthetic-sounding calls, we would expect higher tsik responses. The control condition remained the same as in experiment 1.

Playback condition significantly predicted response scores in linear mixed-effects models (type III ANOVA, F_2,16_ = 5.199, p = 0.018 for phee responses, and F_2,16_ = 5.608, p = 0.014 for tsik responses). Marmosets showed significantly reduced S_phee_ (Fig. 2G, pairwise comparisons in LMM with Tukey adjustment, n=8, df = 19.9, t ratio = -2.772, p = 0.030) and elevated S_tsik_ (Fig. 2H, pairwise comparisons in LMM with Tukey adjustment, n=8, df = 21.3, t ratio = 2.791, p = 0.028) responses during the P-div-sim condition relative to P-ctr. Because identical calls elicited different responses depending on the direction of the playbacks, these effects cannot be explained by absolute acoustic dissimilarity alone. Instead, marmosets tracked how a partner’s calls changed across turns, consistent with models of marmoset vocal accommodation^27^. Calls that diverged over time elicited fewer phee and more tsik responses regardless of absolute acoustic distance, whereas converging calls supported continued affiliative exchange. These results show that marmosets evaluate vocal signals in relation to preceding turns, revealing context-dependent interpretation during communication. We thus can exclude the two alternative explanations for our results (i.e., a mere aversion to synthetic calls, or to calls that are highly dissimilar to their own calls) and find strongest support for the normative interpretation.

### Social reinforcement is partner-specific

In experiment 3, we asked whether the responses to vocal convergence and divergence reflected expectations tied to a specific social relationship or whether they generalized to any conspecific. This distinction is important because vocal accommodation is thought to facilitate social bonding^17,18^, and phee calls in marmosets serve to maintain contact between social partners. Under this view, expectations about vocal alignment should be strongest within established pairs. We therefore repeated experiment 1 with eight adults from four breeding pairs, including two previously tested and two naïve pairs, replacing partner vocalizations with calls from unfamiliar individuals. Subjects were exposed to three interactive antiphonal playback conditions: natural stranger calls (S-ctrl), stranger-like synthetic calls that gradually converged toward (S-con-sim) or diverged from (S-div-dis) the focal individual’s calls (Fig. 2I).

Unlike partner-based playbacks, marmosets showed no significant differences in either phee or tsik responses across the three stranger conditions (Fig. 2J, K). Converging stranger calls did not increase affiliative responses, and diverging stranger calls did not elicit agonistic responses. This indicates that positive and negative reinforcement to acoustic changes is partner-specific.

### Marmosets adjust vocal output to maintain acoustic similarity with partner

Lastly, we asked whether marmosets actively modify their own calls to maintain acoustic similarity during interaction. For each vocal response across experiments 1 to 3, we quantified three measures: acoustic distance to the immediately preceding call that was played back (Prev), mean distance to all playbacks in the pre-treatment baseline (Play), and mean distance to the individual’s own pre-treatment baseline calls (Self). Together, these measures can distinguish convergence toward playbacks, divergence from them, or no systematic change (see predictions across stimulus conditions in Fig. 3).

**Fig 3.**
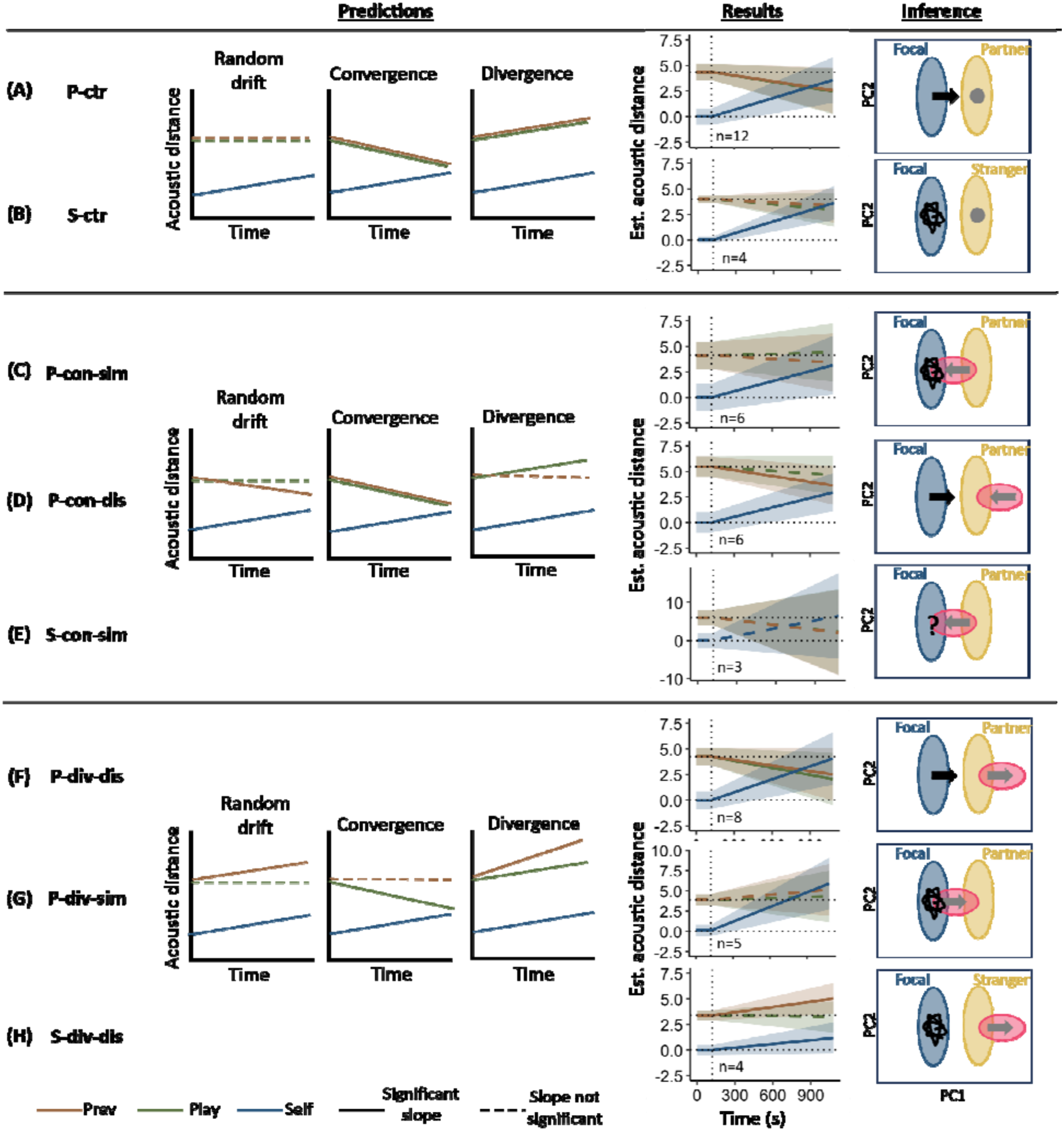
Marmosets actively adjust vocal output to maintain similarity with social partners. The figure shows model estimates of changes in acoustic distance across playback sessions for all stimulus conditions in experiments 1 to 3. Acoustic distances were quantified relative to three reference measures: the immediately preceding playback call (Prev), the set of playback call presented during the initial baseline period (Play), and the focal individual’s own calls produced during the initial baseline period (Self). Lines in the predictions panel illustrate the expected trajectories under three hypothetical response strategies: random drift, vocal convergence, or vocal divergence. Lines in the results indicate Theil-Sen trends across time and shaded regions denote 95% confidence intervals. Marmosets converged toward partner-derived playback stimuli in the partner control **(A)** condition supporting the results that convergence is the norm. Marmosets also converged to the playbacks in the simulated divergence **(F)** and reversed convergence **(D)** conditions, indicating active compensation for increased acoustic dissimilarity. In contrast, random drift best explained the results during all other conditions **(B**,**C**,**G**,**H)** while the number of responses to the stranger convergence playbacks were insufficient to make any inferences **(E)**. Abbreviations: Prev, acoustic distance between each focal response and the immediately preceding playback; Play, mean acoustic distance between each focal response and playback calls from the initial baseline period; Self, mean acoustic distance between each focal response and the focal individual’s own calls from the initial baseline period.

To capture both the direction and the magnitude of the rate of change over time, we estimated trends in each measure using non-parametric robust Theil-Sen slopes (as opposed to parametric least-squares regression). In experiment 1, marmosets showed clear convergence toward playback stimuli in the P-ctr condition, consistent with short-term vocal accommodation (Fig. 3A, n = 12, 95% C.I. for Prev slopes = [-0.00336, -0.00040] and for Play slopes = [-0.00346, - 0.00050]). Notably, convergence was also observed in conditions where playbacks became more dissimilar to the partners’ calls than the focal’s own calls, that is, P-div-dis (Fig. 3D, n = 8, 95% C.I. for Prev slopes = [-0.00358, -0.00003] and for Play slopes = [-0.00406, -0.00051]) and P-con-dis (Fig. 3F, n = 6, 95% C.I. for Prev slopes = [-0.00282, -0.00098] and for Play slopes = [-0.00177, 0.00007]). In these cases, individuals adjusted their vocal output to reduce the acoustic difference. In all other conditions, marmosets drifted away from their pre-baseline vocalizations without significant convergence or divergence to the playbacks (Fig. 3B, C, G, H). Phee responses in the stranger convergence condition were too few for meaningful inference (S-consim, Fig. 3E).

The above pattern indicates that marmosets actively regulate vocal output to maintain acoustic similarity with the partner. Specifically, in situations of high acoustic distance with their partner, marmosets converge to counteract the dissimilarity. In line with the probability-to-reply findings, such compensatory vocal adjustments were not observed with strangers, suggesting that this process is partner-specific.

## Discussion

Our results show that common marmosets expect specific patterns of vocal accommodation from their bond partners, and engage in *positive social reinforcement* (“approval”) when these expectations are met and *negative social reinforcement* (“disapproval”) when they are violated. Marmosets increased affiliative phee responses to calls that became progressively more similar to their own, consistent with positive social feedback (Fig. 2D). In contrast, calls that diverged from the focal individual reduced affiliative responses (Fig. 2G) and elicited aversive tsik vocalizations, indicating negative social feedback (Fig. 2E, 2H). Importantly, tsiks stopped when calls reverted to baseline (Fig. S2), suggesting a communicative repair function^28^. These responses were specific to established social partners and could not be explained by low-level alternative explanations (Fig. 2J, 2K). Responses to acoustic changes in partners, but not strangers, were accompanied by active vocal adjustments that reduced acoustic differences (Fig. 3). Together, these findings demonstrate that marmosets track changes in a partner’s calls^27^, react negatively to partner divergence, and regulate their own vocal output to maintain acoustic similarity.

These properties align with core features of social norms^6–9^. Previous work has shown that vocal accommodation in marmosets is expressed *consistently* across individuals^15^ with convergence as the dominant pattern, and divergence occurring only rarely, suggesting *conformity*^16^. Our results add a critical component by demonstrating that this behavior is *socially reinforced*. Vocal accommodation in marmosets therefore satisfies key criteria for norm-like behavior, making it a strong candidate for a socially maintained vocal norm in a nonhuman primate.

Evidence for socially shared conventions in nonhuman primates comes primarily from studies of great ape behavior and gesture, as well as group-level conformity in tool use and food preferences ^29–36^. These studies show that nonhuman primates can learn and adopt locally shared behavioral practices, and that great apes may share expectations about how some gestures are used and interpreted. However, evidence for a normative organization of such regularities remains limited. In particular, there is little evidence that individuals actively approve of expected behavior or disapprove of violations, a key feature of social norms in humans^9,14^. Our findings therefore extend beyond the documentation of shared behavioral conventions by identifying affiliative and aversive responses to conformity and deviation in a communicative behavior, providing evidence supporting a social-vocal norm in a nonhuman primate.

Understanding the evolutionary origins of communicative norms in humans requires identifying precursor systems. Vocal accommodation in marmosets provides a model for studying how shared expectations about signal form emerge, are maintained, and influence interaction, supporting the idea that human language builds on basic mechanisms of socially regulated communication^18,27,37,38^. At the same time, our evidence is currently limited to dyadic interactions, whereas human linguistic norms are sustained at the group level and extend beyond individuals directly involved in an interaction. Determining whether marmoset communicative norms are similarly shared and enforced by third parties could clarify whether such phenomena represent a precursor to the group-level normativity characterizing human language.

## Acknowledgments

We are thankful to Carel P. van Schaik and Hans-Johann Glock for the discussions on the manuscript and Rahel K. Brügger for her inputs on the statistical analyses.

## Author contributions

Conceptualization: NP, JMB

Methodology: NP, JMB, MM

Investigation: NP

Visualization: NP

Funding acquisition: JMB

Project administration: JMB

Supervision: JMB

Writing – original draft: NP

Writing – review & editing: JMB, MM

## Data, code, and materials availability

All data and code used in the analysis are available at https://doi.org/10.5281/zenodo.23496574. The dataset will be made public upon publication.

