## Extended figures for "Norm-like vocal behavior in a nonhuman primate"

**Affiliations:**

**The PDF file includes:**

Figs. S1 to S2

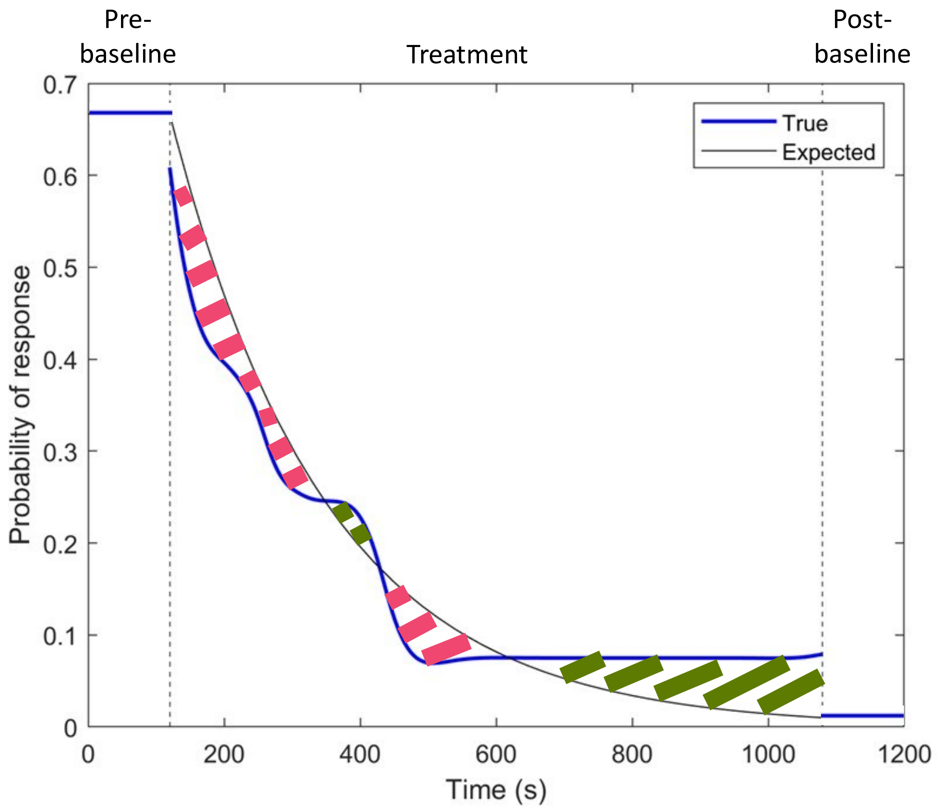


**Fig. S1. Illustration of the procedure used to quantify vocal response scores.** Example session showing the expected response probability trajectory (black) and the observed true response probability calculated using a sliding 2-minute window (blue). The mean response probability during the first and final 2 minutes of the session defined the pre- and post- baselines, respectively. These baseline values were used to parameterize an expected exponential decline in response probability across the treatment period. The response score $S_{call}$ was calculated as the time-integrated difference between the observed and expected response probabilities from the end of the pre-baseline period (120 s) to the start of the post-baseline period (1080 s). Green shaded regions indicate periods where the true response probability exceeded the expected trajectory and contributed positively to the response score, whereas pink shaded regions indicate periods where the observed response probability fell below expectation and contributed negatively. Positive response scores therefore reflect greater-than-expected responsiveness to playback stimuli, while negative scores reflect reduced responsiveness.

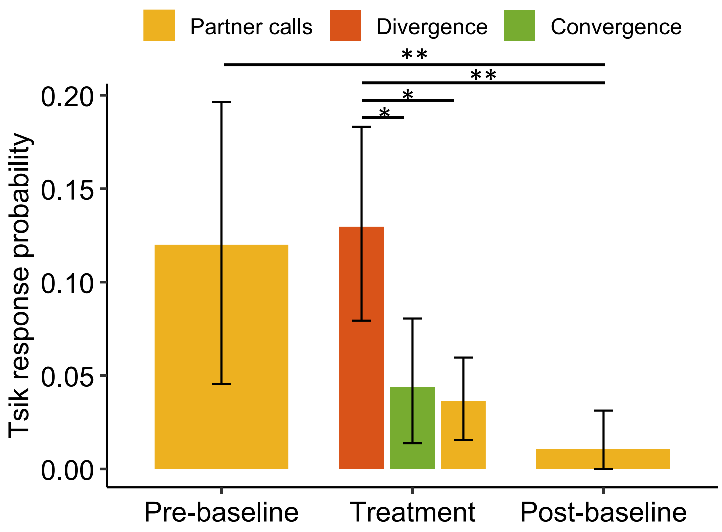


**Fig. S2. Agonistic tsik responses are elevated during simulated vocal divergence.** Mean tsik response probabilities during the pre-baseline, treatment, and post-baseline periods. Pre- and post-baseline values are shown for the divergence condition only, whereas treatment-period values are shown for partner control, convergence, and divergence conditions. Marmosets produced significantly more tsik calls during simulated vocal divergence than during partner-control and convergence playbacks. Tsik response probabilities reduced drastically from the treatment to post-baseline period for the divergence condition. Bars represent means and error-bars indicate 95% bootstrapped confidence intervals. Asterisks denote post hoc comparisons in the model with Holm correction (*$p < 0.05$, **$p < 0.01$).
