## Supplementary tables and videos for "Norm-like vocal behavior in a nonhuman primate"

**Affiliations:**

**The PDF file includes:**

Tables S1 to S2
Legends for Movies S1 to S2

**Table S1. Model selection and significance testing for linear mixed-effects models of phee response scores in experiment 1.** Type III ANOVA results for the model containing playback condition as the sole fixed effect are shown in the first row. The second row reports the likelihood-ratio test (LRT) comparing the base model to a null model containing only random effects, assessing the overall contribution of playback condition. Subsequent rows report LRTs evaluating whether the addition of age, sex, presentation order, or their interactions with playback condition significantly improved model fit compared to the condition-only model. ΔAIC values indicate the change in Akaike Information Criterion relative to the simpler model in each comparison. Negative values indicate improved fit of the more complex model. All models included individual identity nested within pair identity as random effects.

| Term | Chisq | Df | P_value | ΔAIC |
|---|---|---|---|---|
| **Condition (condition-only model)** | **—** | **2** | **1.27E-06** | **—** |
| **Condition (vs null model)** | **27.446** | **2** | **1.10E-06** | **-23.446** |
| Condition + Age | 0.946 | 1 | 0.331 | 1.054 |
| Condition + Sex | 1.038 | 1 | 0.308 | 0.962 |
| Condition + Order | 1.496 | 4 | 0.827 | 6.504 |
| Condition x Age | 3.433 | 2 | 0.18 | 0.567 |
| Condition x Sex | 0.587 | 2 | 0.746 | 3.413 |
| Condition x Order | 5.234 | 8 | 0.732 | 10.766 |

**Table S2. Evaluation of fixed effects in linear mixed-effects models of tsik response scores in experiment 1.** The first row reports the significance of playback condition in the model which included condition as the only fixed effect. The second row compares this model to a null model containing only random effects using a likelihood-ratio test (LRT), providing a measure of the overall contribution of playback condition. Subsequent rows show LRTs testing whether the inclusion of age, sex, presentation order, or their interactions with playback condition improved model fit compared to the condition-only model. Changes in Akaike Information Criterion (ΔAIC) are reported for each model comparison. All models included individual identity nested within pair identity as random effects.

| Term | Chisq | Df | P_value | ΔAIC |
|---|---|---|---|---|
| **Condition (condition-only model)** | **—** | **2** | **0.000106** | **—** |
| **Condition (vs null model)** | **18.311** | **2** | **0.000106** | **-14.311** |
| Condition + Age | 0.019 | 1 | 0.889 | 1.981 |
| Condition + Sex | 0 | 1 | 0.989 | 2 |
| Condition + Order | 2.782 | 4 | 0.595 | 5.218 |
| Condition x Age | 0.926 | 2 | 0.629 | 3.074 |
| Condition x Sex | 0.073 | 2 | 0.964 | 3.927 |
| Condition x Order | 4.27 | 8 | 0.832 | 11.73 |

**Movie S1. Example segment of a partner control trial showing the automated closed-loop acoustic playback paradigm.** The video focuses on the cage where the focal is present, while the speaker is located behind the white opaque visual barrier to the right of the cage. The video has been sped up by 3X.

**Movie S2. An example divergence trial (P-div-dis).** The video focuses on several timepoints within the pre-baseline, divergence and post-baseline segments where it is sped up by 3X and both phee and tsik responses from the focal can be heard. The video is sped up by 15X for the remainder of the trial. The bar at the bottom of the video, adapted from Fig. 2C, shows the current stage in the timeline of the trial.
